# Inhibition of mTORC1 signaling in aged rats counteracts the decline in muscle mass and reverses multiple parameters of muscle signaling associated with sarcopenia

**DOI:** 10.1101/591891

**Authors:** Giselle A. Joseph, Sharon Wang, Weihua Zhou, Garrett Kimble, Herman Tse, John Eash, Tea Shavlakadze, David J. Glass

## Abstract

There is a lack of pharmacological interventions available for sarcopenia, a progressive age-associated loss of muscle mass, leading to a decline in mobility and quality of life. We found mTORC1 (mammalian target of rapamycin complex 1), a well-established critical positive modulator of mass, to be hyperactivated in sarcopenic muscle. Furthermore, inhibition of the mTORC1 pathway counteracted sarcopenia as determined by observing an increase in muscle mass and fiber type cross sectional area, surprising because mTORC1 signaling has been shown to be required for muscle mass gains in some settings. Additionally, several genes related to senescence were downregulated, while gene expression indicators of neuromuscular junction denervation were diminished using a low dose of a rapalog. Therefore mTORC1 inhibition may delay the progression of sarcopenia by directly and indirectly modulating multiple age-associated pathways, implicating mTORC1 as a therapeutic target to treat sarcopenia.

## Introduction

Skeletal muscle size is physiologically regulated by load and activity, and can decrease when load is reduced. Muscle also atrophies, or decreases in size, in pathological conditions such as cancer, immobilization and denervation (1). One setting where muscle mass and function are diminished is old age. This loss of muscle is called sarcopenia, and it is associated with a decrease in the ability to move, leading to morbidity and ultimately mortality (2); indeed, a decrease in walking speed is one of the strongest predictors of mortality in humans, and this finding is associated with sarcopenia (3, 4). In addition to frailty and sarcopenia, aging of course affects every tissue system and greatly increases susceptibility to other serious diseases and co-morbidities, such as cancer, heart failure, chronic kidney disease, loss of vision, dementia and Alzheimer’s disease (1, 5, 6).

Experimental data strongly suggest the coordinated regulation of aging by distinct molecular pathways (7); modulation of these pathways can counteract several age-related diseases and co-morbidities, and prolong life (7-10). Of these signaling pathways, genetic or pharmacological inhibition of the mammalian target of rapamycin (mTORC1) is thus far the best-validated intervention to delay age-related pathophysiological changes (11). For instance, the use of an mTORC1 inhibitor, rapamycin, even when administered at later stages in life, has been shown to extend lifespan in mice (12-15). Pharmalogical agents related to rapamycin are called “rapalogs”. Use of a rapalog for aging-like indications has recently been translated to human beings, where it was shown to improve responses to vaccinations in the elderly, coincident with decreasing signs of immune-senescence (16). The low dose rapalog treatment used in the human study was reverse-translated to rats, where it was shown that intervention late in life could prevent signs of age-related kidney pathology (17). However, there has always been concern about the potential effects of rapamycin and rapalogs on skeletal muscle. For example, inhibition of the mTORC1 pathway was shown to entirely block responses to compensatory hypertrophy in mice (18). This certainly gave the impression that activation of mTORC1 signaling was desireable for the maintenance of muscle mass. Most recently it was shown that rapamcyin treatment inhibited muscle mass increase caused by myostatin loss (19). Thus it seemed reasonable that inhibition of the pathway was not desireable in settings of muscle loss (1, 18, 20).

As to the pathway, Akt induces protein synthes in part by activation of mTORC1 signaling (18, 21). mTOR exists in the distinct complexes, mTORC1 and mTORC2. mTORC1 is characterized by the presence of RAPTOR (regulatory-associated protein of mTOR) (22), while TORC2 binds to RICTOR (rapamycin-insensitive partner of mTOR) (23, 24). The mTORC1 complex induces downstream signaling responsible for protein synthesis through phosphorylation and activation of S6 Kinase 1 (S6K1), and via inhibition of 4E-BP1 (24, 25), and is sensitive to inhibition by rapamycin and rapalogs. In addition to the anabolic function, Akt also limits muscle protein degradation and atrophy by phosphorylating and thereby inhibiting the FOXO (also known as Forkhead) family of transcription factors. Activation of FOXO3 is sufficient to induce atrophy (26, 27); transgenic expression of FOXO1 also lead to an atrophic phenotype (28, 29). Dephosphorylated FOXO1 and FOXO3 proteins translocate to the nucleus where they induce transcription (30), upregulating the expression of the muscle-atrophy associated E3 ligases, muscle RING finger 1 (MuRF1) and muscle atrophy F-box (MAFbx)/Atrogin-1 (31-33). Both MuRF1 and MAFbx/Atrogin-1 are specifically upregulated in atrophic conditions (34, 35), and target proteins that are critical for muscle structure and protein synthesis for degradation, thereby inducing muscle loss (36-40).

mTORC1 inhibition has been widely suggested as a way to improve function in the elderly in various tissues. However, its potential as a therapeutic intervention for the treatment of sarcopenia has not been considered. Upon examination, we were surprised to learn that mTORC1 signaling is upregulated rather than downregulated coincident with signs of sarcopenia in rats, We therefore explored the effects of rapalog treatment in this setting. The results demonstrate that the inhibition of mTORC1 is helpful in preventing pathological changes related to sarcopenia.

## Results

### Increased activation of the mTORC1 pathway with age

Given prior reports that mTORC1 inhibition was helpful to treat a variety of age-related disorders, but also the data that mTORC1 activation is required for muscle hypertrophy, we conducted a time course analysis of the mTORC1 pathway to get a full scope of how its activity changes with age. In laboratory settings, Sprague Dawley rats have an average lifespan of up to 2.5 to 3 years (41). In our study, male rats ranging from 6-months to 27-months were used. Protein lysates from gastrocnemius muscles were probed for the downstream effector of mTORC1, phosphorylated ribosomal protein S6 (rpS6), as a determinant of pathway activity. Basal (6 hours fasted) levels of phosphorylated rpS6 gradually increased as the rats aged, with a substantial increase of about 10-fold in the oldest animals aged 27-months compared with 6-months. (Fig. 1A, B). The age-related increase in mTORC1 signaling coincided with a decrease in muscle mass. At 21-months gastrocnemius muscle weights declined and progressively atrophied at each later time point (Fig. 1C). Though muscle loss at this age is not a surprise, the coincidence of this loss with mTORC1 activation was quite unexpected, given that it favors muscle growth and hypertrophy.

**Figure 1.**
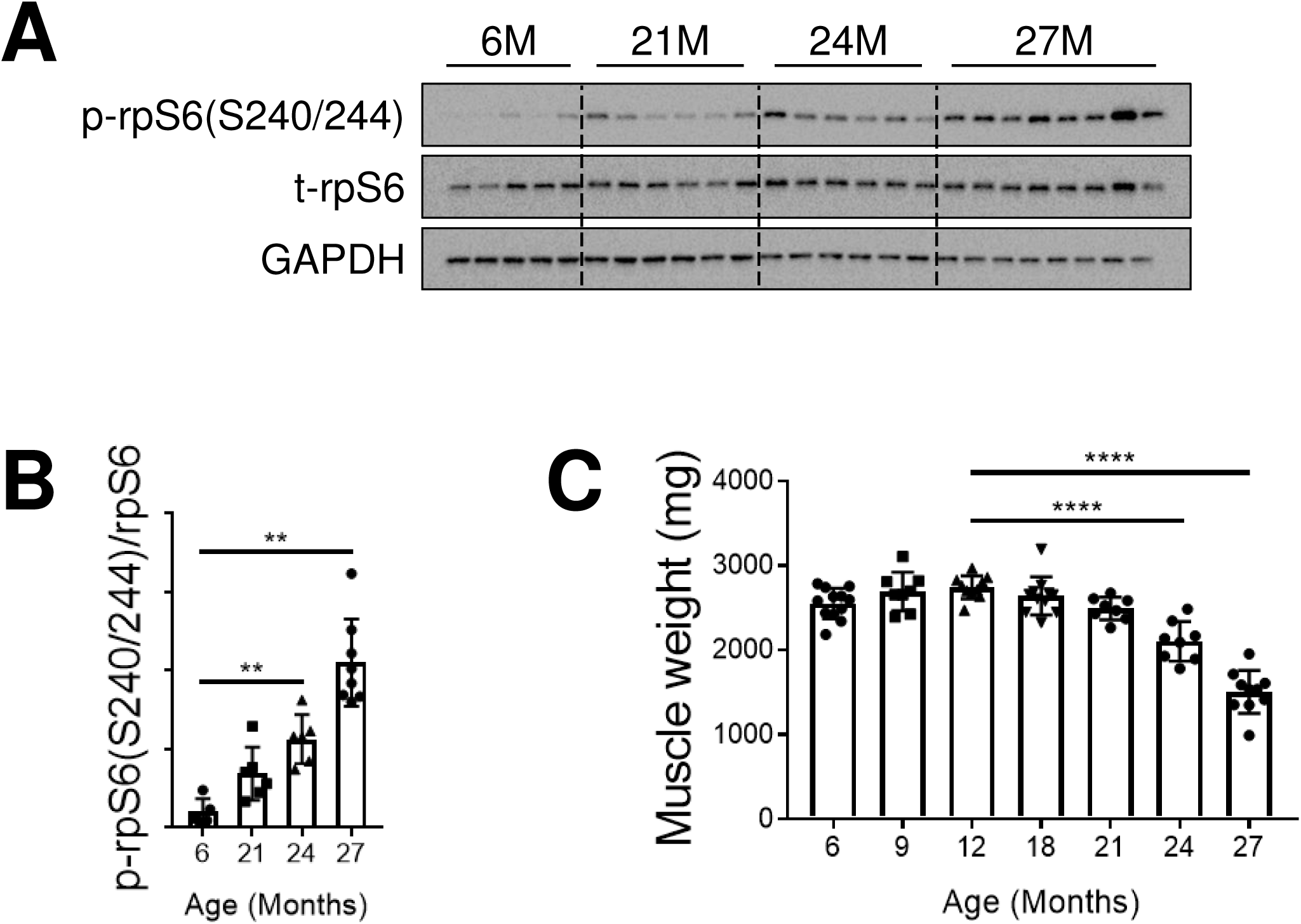
mTORC1 signaling is hyper-activated in sarcopenic skeletal muscle. (A) Immunoblots for phosphorylated (p) and total (t) protein for rpS6 in gastrocnemius muscles of rats aged 6 (n = 5), 21 (n = 6), 24 (n = 6) and 27 (n = 8) months. Glyceraldehyde-3-phosphate dehydrogenase (GAPDH) is shown as a loading control. (B) p-rpS6(S240/244) protein amounts were quantified relative to their respective total rpS6 protein amounts by densitometry. (C) Gastrocnemius muscle weights in rats aged 6 (n = 12), 9 (n = 8), 12 (n = 10), 18 (n = 11), 21 (n = 8), 24 (n = 8) and 27 (n = 10) months. Data are mean ± standard deviation of the mean. Statistical significance was determined by a one way ANOVA followed by Dunnett’s multiple comparison tests. Means from all groups were compared to the mean of 12-month old animals. Asterisk (*) denotes significance at **P< 0.01; ***P< 0.001 and ****P< 0.0001. Y-axis in (B) represents arbitrary units, and in (C), milligrams (mg).

### Skeletal muscle mass and quality is improved in sarcopenic rats treated with the rapalog RAD001

Experimental evidence shows that the use of rapalogs as therapeutic agents is beneficial in extending lifespan and counteracting age-related morbidities in humans and other evolutionarily diverse species (reviewed in (42)). We sought to determine whether rapalog treatment could counter the pathophysiological changes associated with sarcopenia. Aging rats display signs of sarcopenia beginning at 18-months (43). In the present study, aged rats (22-months) were dosed daily with either vehicle or 0.15mg/kg RAD001 for 6-weeks. This dose of RAD001 is equivalent to a clinical dose of 0.5mg in humans, ensuring therapeutic relevance (16). Vehicle treated young adult rats (7-months) served as a comparative baseline for aging effects. At the end of the treatment, aged and young adult rats were 24-months and 9-months old respectively, and will be referred to as such.

To determine if we were able to ameliorate age-related muscle loss with rapalog treatment, we measured the wet weights of the tibialis anterior (TA), plantaris, gastrocnemius, and soleus muscles. Consistent with previous data, all muscles, except for the soleus muscle from 24-month old vehicle treated rats had considerably reduced mass compared to 9-month old rats (Fig. 2A). RAD001 treatment did not lead to further atrophy in any of these muscles. On the contrary, rapalog appeared to be protective for aged animals, and reduced extensive muscle mass loss. Plantaris and TA muscles showed a surprising increase in mass, with the TA muscle being significantly increased compared to vehicle treated animals (Fig. 2A). Our data provide strong evidence that when administered to sarcopenic rats, low dose rapalog treatment is not detrimental to muscle mass. Rather, it allows for animals to maintain or gain muscle.

**Figure 2.**
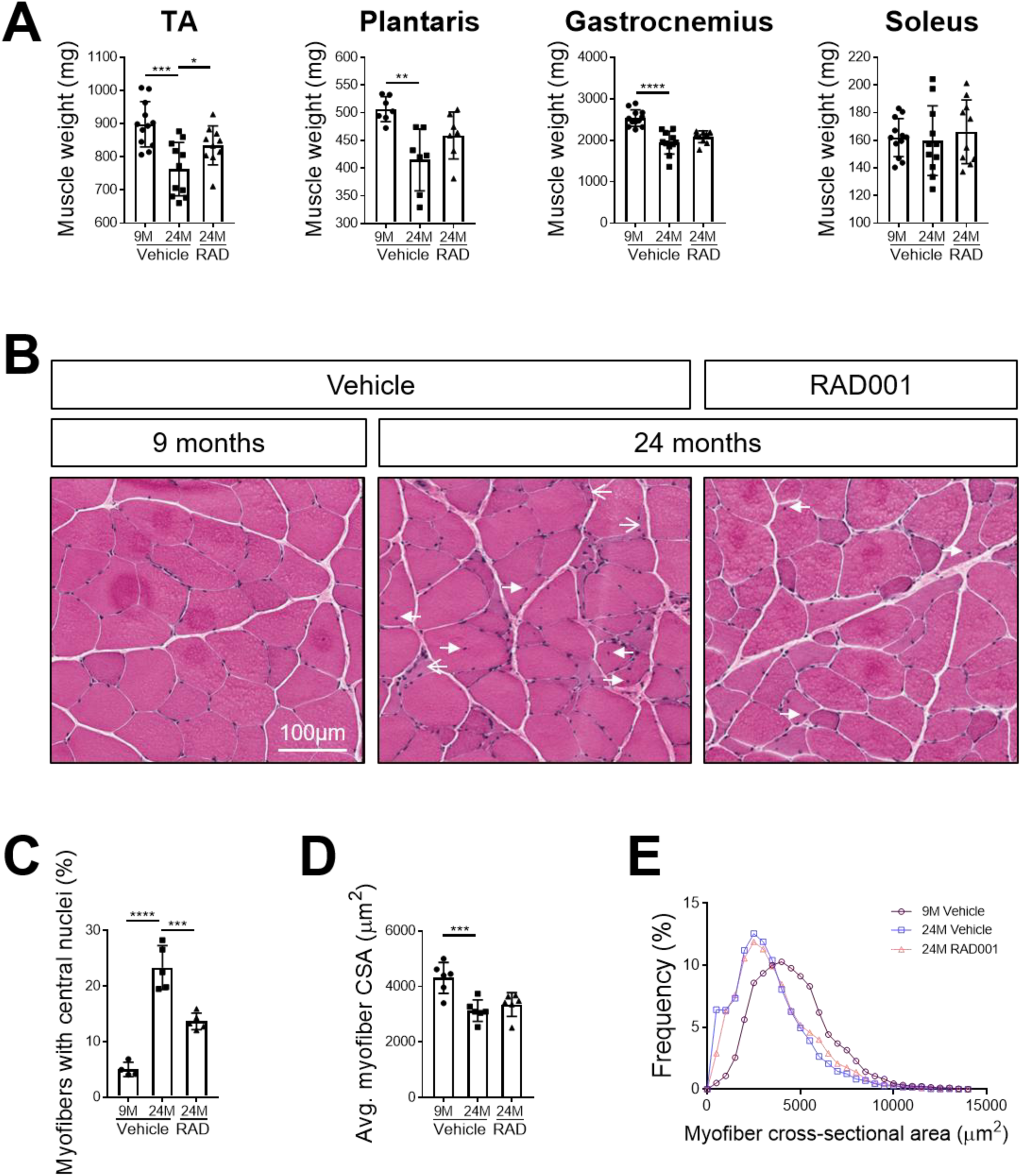
Skeletal muscle mass and quality is improved with RAD001 treatment. (A) Weights of tibialis anterior (TA), plantaris, gastrocnemius and soleus muscles from 9- and 24-month old rats treated with vehicle and 24-month old rats treated with RAD001 (n = 7-12 animals per group). Y-axes represent weight units in milligrams (mg). (B) Representative images of transverse sections of plantaris muscles stained with H&E from 9- and 24-month old rats treated with vehicle (n = 4 and n = 5 animals per group respectively) and 24 month old rats treated with RAD001 (n = 5 animals). Open arrows depict mis-shaped, flattened myofibers. Closed arrows show central nuclei. Scale bar is 100 μm. (C) Quantification of myofibers withcentral nuclei in plantaris muscles. Myofibers with central nuclei are shown as a percentage of total myofibers (n> 1200 myofibers assessed per animal). (D) Average myofiber cross-sectional area (CSA) in plantaris muscles (n > 1200 myofibers assessed per animal). Y-axes represent area units in microns squared (µ^2^). (E) Histogram depicting the distribution of myofiber cross-sectional areas from data shown in (D). Myofiber cross-sectional area frequencies are shown as a percentage of total myofibers in the given treatment group. All other data are mean ± standard deviation of the mean. Asterisk (*) denotes significance at *p< 0.05; **p< 0.01; ***p< 0.001 and ****p< 0.0001.

Changes in muscle mass often reflect morphological alterations in tissue. We performed histological analysis on H&E stained plantaris muscle cross-sections. Tissue from 9-month old rats had normal morphology, typical of healthy muscle (Fig. 2B). In contrast, we detected several indicators of distressed muscle in aged animals that received only vehicle. A high proportion of fibers had a smaller cross-sectional area, a phenotype associated with muscle atrophy (Fig. 2D, E). Moreover, about 23% of myofibers from vehicle treated 24-month old muscles presented with central nuclei, indicative of prior degeneration and ongoing regeneration (Fig. 2B, C). There was a striking reduction in the number of myofibers with central nuclei in 24-month old plantaris muscles treated with RAD001 compared with aged matched muscles treated with vehicle (Fig. 2B, C). In addition, consistent with the observed trend of increased plantaris muscle mass in RAD001 treated rats, the average myofiber cross-sectional area tended to increase (Fig. 2D) - the most obvious change was a reduced frequency of very small, mis-shaped atrophic myofibers (Fig. 2E). Taken together, these data show that a low dose rapalog treatment for 6 weeks can counteract age-related morpho-pathological changes in sarcopenic skeletal muscle - especially signs of degeneration requiring regeneration, as measured by the presence of central nuclei.

Chronic activation of the mTORC1 pathway by muscle-specific deletion of *Tsc1*, a negative regulator of mTORC1, has been shown to cause a late-onset myopathy, with muscle atrophy in young adult mice (44). Inhibition of mTORC1 activity using rapamycin was able to reverse the observed pathological changes and normalize muscle mass in these animals (44). Despite evidence of similarly sustained mTORC1 signaling in aged muscle, its inhibition has not yet been studied in the context of sarcopenia. Given the positive phenotypic changes in aged RAD001-treated muscle, we assessed perturbation of mTORC1 pathway activity. Western blot analyses confirmed that RAD001 treatment significantly reduced phosphorylation of S6K1, a downstream target of mTORC1, in the muscles of old rats (Fig. 3A, B; Fig. S1). Phosphorylation of rpS6, a direct downstream target of S6K1 was also reduced with RAD001 treatment (Fig. 3A, C; Fig. S1). These data confirm that the relatively low dose of the rapalog used in the present study was sufficient to inhibit mTORC1 signaling in aged skeletal muscle.

**Figure 3.**
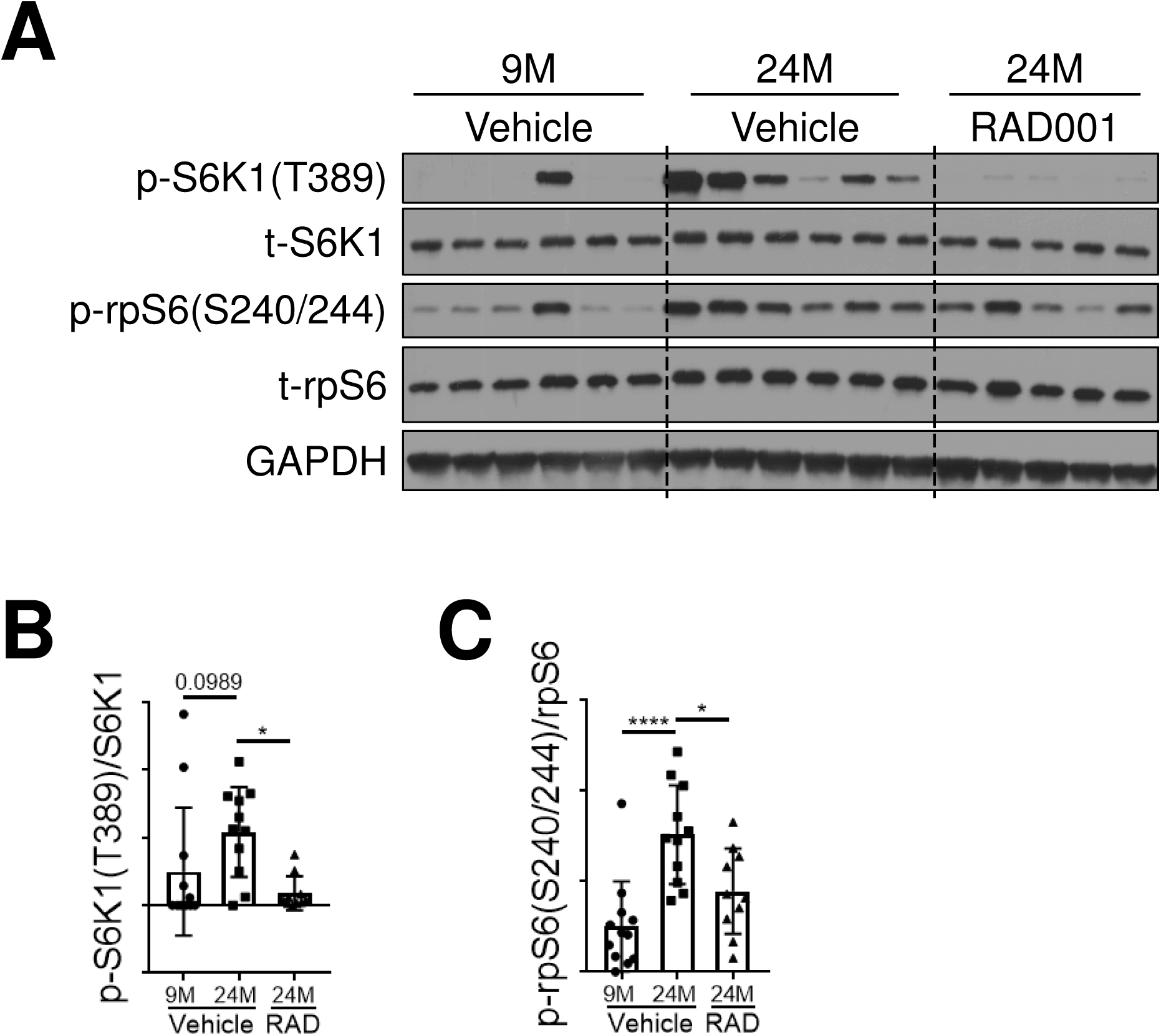
Confirmed mTORC1 inhibition following rapalog treatment. (A) Representative immunoblots for phosphorylated (p) and total (t) protein for S6K1 and rpS6 in tibialis anterior muscles of 9- and 24-month old rats treated with vehicle and 24-month old rats treated with RAD001. Glyceraldehyde-3-phosphate dehydrogenase (GAPDH) is shown as a loading control. (B) p-S6K1(T389) and (C) p-rpS6(S240/244) protein amounts were quantified relative to their respective total S6K1 and rpS6 protein amounts by densitometry (n = 10 – 12 animals per group). Data are mean ± standard deviation of the mean. Asterisk (*) denotes significance at ***p< 0.001.Y-axes represent arbitrary units.

### mTORC1 inhibition reverses molecular changes associated with sarcopenia

We previously reported on age-related gene expression changes that help to demonstrate the molecular pathogenesis of sarcopenia (43). These data revealed the transcriptional upregulation of several pathways, including pathways related to innate inflammation and senescence, cellular processes modulated by mTORC1. Because RAD001-treated animals displayed a remarkable sparing of muscle morho-pathology and mass with mTORC1 inhibition, we sought to determine the molecular changes that could account for this.

The E3 ligase MuRF1 is an important regulator of atrophy (1, 34). MuRF1 gene expression was analyzed in young and old muscles treated with vehicle, and in old muscles treated with RAD001 (Fig. 4A). Old muscles treated with vehicle had significantly higher level of MuRF1 mRNA compared with young muscles (Fig. 4A). RAD001 treatment reduced MuRF1 gene expression in old muscles (Fig. 4A). MaFbx, like MuRF1, is an E3 ubiquitin ligase that is transcriptionally upregulated under atrophic conditions (34). There was a significant increase in MaFbx expression in muscle from 24-month vehicle-treated animals compared to young controls. However, its expression was not impacted by RAD001 treatment (Fig. 4A).

**Figure 4.**
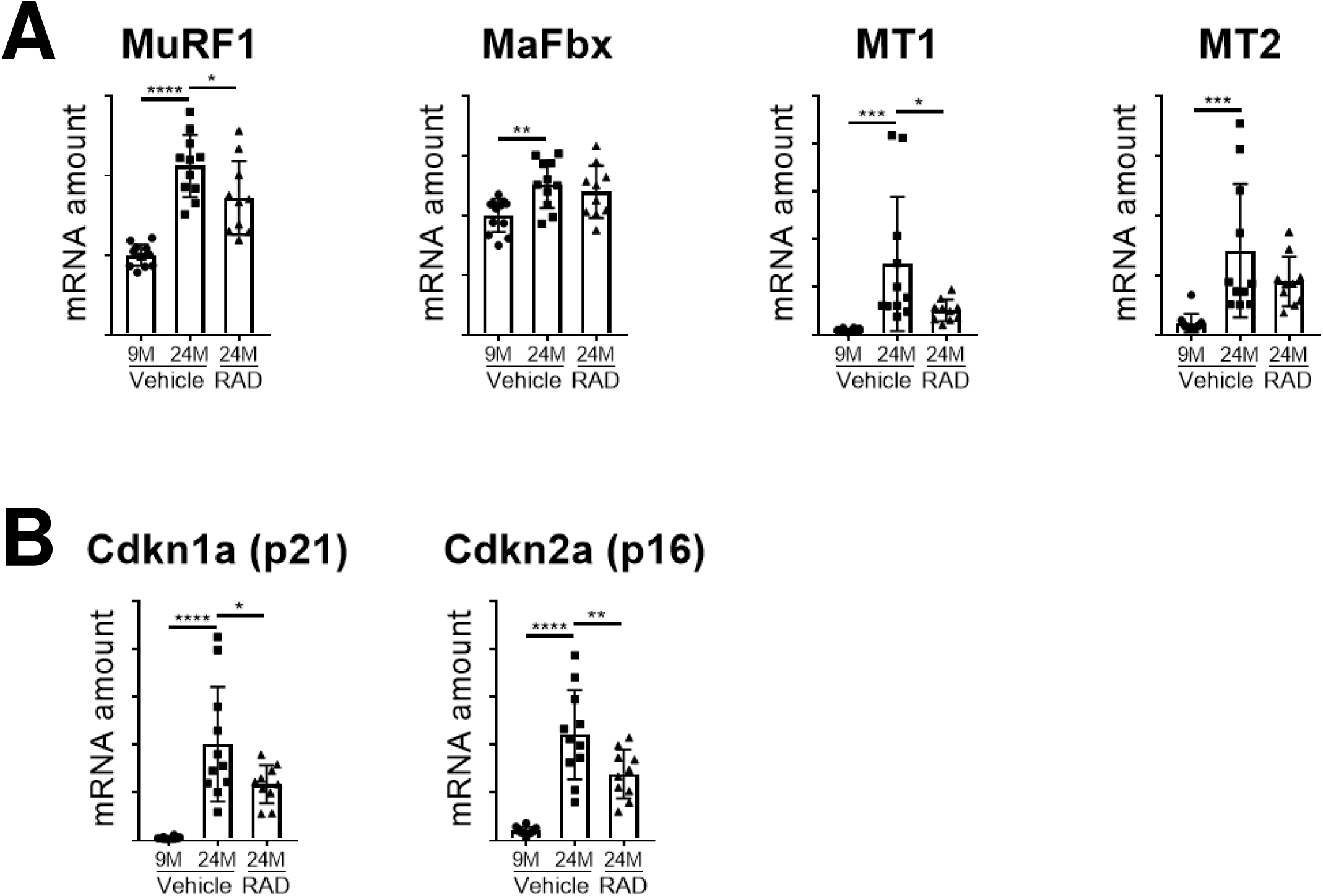
mTORC1 inhibition blunts molecular changes associated with sarcopenia. (A) mRNA amounts for MuRF1, MaFbx, MT1 and MT2, and (B) Cdkn1a and Cdkn2a in tibialis anterior muscles of 9- and 24- month old rats treated with vehicle and 24-month old rats treated with RAD001 (n = 10 – 12 animals per group). mRNA amounts were standardized to a geometric mean of TBP and Vps26a, used as reference genes (A and B). Data are mean ± standard deviation of the mean. Asterisk (*) denotes significance at *p< 0.05; **p< 0.01; ***p< 0.001; ****p< 0.0001. Y-axes represent arbitrary units.

Additionally, expression of the metallothioneins, MT1 and MT2, increases during atrophy and is elevated in sarcopenic muscle (45). Genetic silencing of these genes promotes muscle hypertrophy *in vivo* (45). Interestingly, RAD001 treatment suppressed the MT1 gene expression level in old muscles, while levels of MT2 though variable within either treatment group, remained unchanged between groups (Fig. 4A).Together with the observed increase in muscle mass (Fig. 2A), these data demonstrate that mTORC1 inhibition by RAD001 can prevent further muscle loss by suppressing expression of atrophy markers.

The onset of senescence with age is associated with the inability to efficiently repair and recover muscle, a contributing factor to the progressive decline in muscle mass in sarcopenia. Cell cycle proteins Cdkn1a (p21) and Cdkn2a (p16) are known cellular senescence markers that are upregulated with age in several tissues, including skeletal muscle (43, 46, 47). Relative to 9-month old rats, Cdkn1a (p21) and Cdkn2a (p16) are both highly expressed at the mRNA level in muscle from aged vehicle treated rats (Fig. 4B). RAD001 significantly reduced Cdkn1a and Cdkn2a mRNA levels in 24-month old muscles compared to age-matched vehicle treated controls (Fig. 4B).

### RAD001 treatment protects from age-associated signs of denervation

Along with the deterioration of muscle tissue, the breakdown of the neuromuscular junction (NMJ) also contributes to muscle weakness; abrogation of the NMJ is associated with aging in rodents (48-50). Previous work identified the transcriptional perturbation of several genes associated with functional denervation and the loss of motor neurons in the rat sarcopenic muscle (43). We therefore investigated whether mTORC1 inhibition could reverse these transcriptional changes. The expression levels of a panel of select genes, Chrna1, Chrne, MuSK, Myogenin and Gadd45a, known to be markers of functional denervation (43), were determined by RT-qPCR. In agreement with our previous observations, all of these genes were significantly upregulated in muscles from vehicle treated 24-month old animals compared to 9-month old controls (Fig. 5). Interestingly, treatment of aged animals with RAD001 reduced the transcriptional upregulation of these denervation-associated gene markers in relation to their vehicle treated age-matched counterparts (Fig. 5). These data suggest that suppressing the mTORC1 pathway in aged animals could be protective against age-associated denervation.

**Figure 5.**
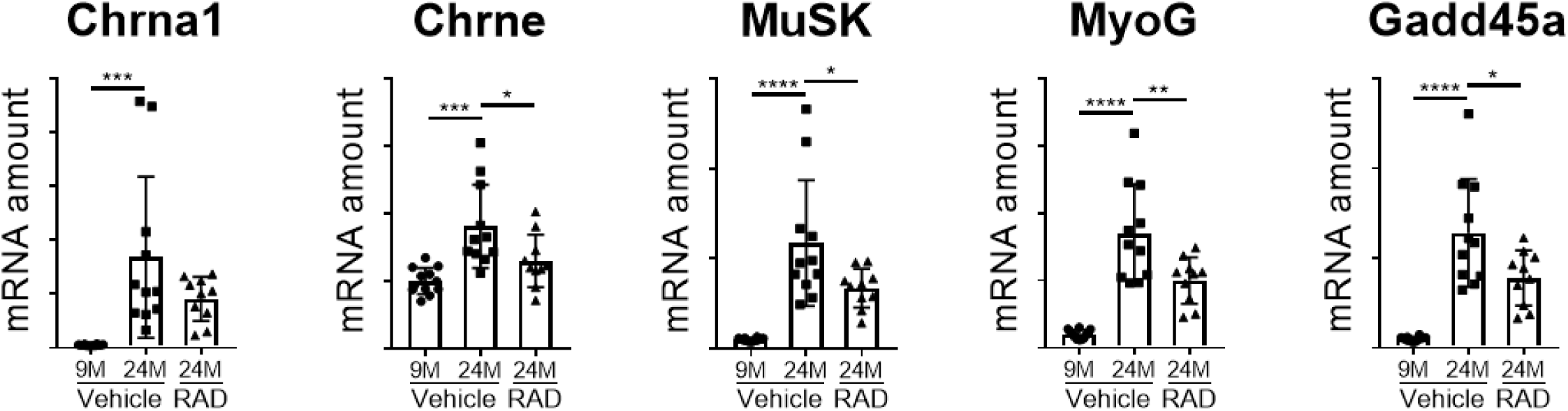
mRNA expression of denervation markers are reduced by rapalog treatment. (A) mRNA amounts for Chrna1, Chrne, Musk, MyoG and Gadd45a in tibialis anterior muscles of 9- and 24-month old rats treated with vehicle and 24-month old rats treated with RAD001 (n = 10 – 12 animals per group). mRNA amounts were standardized to a geometric mean of TBP and Vps26a, used as reference genes. Data are mean ± standard deviation of the mean. Asterisk (*) denotes significance at *p< 0.05; **p< 0.01; ***p< 0.001; ****p< 0.0001. Y-axes represent arbitrary units.

## Discussion

Age-associated diseases comprise many of the most serious conditions afflicting human beings: sarcopenia and frailty, cancer, heart disease, Alzheimer’s disease, and chronic kidney disease. The mTORC1 inhibitor rapamycin and its analogs (rapalogs) have been shown to extend lifespan (12-15) and delay many of these age-related conditions (9-11). These findings have even been extended to human beings, where a rapalog reversed immune-senescence and increased responses to vaccines that normally decline with age in the elderly (16). One area which caused some concern when it came to giving mTORC1 inhibitors to aged subjects was skeletal muscle, since mTORC1 activation mediates protein synthesis (51) and mTORC1 inhibition blocks load-dependent hypertrophy (52). However, when we examined mTORC1 signaling in skeletal muscles in rats, at ages where sarcopenia occurs (43), we were surprised to see that signaling was increased rather than decreased - there was an age-related increase in the phosphorylation of rpS6, a readout of mTORC1 activity. Coincident with elevated mTORC1 signaling, there was a progressive decrease in skeletal muscle mass. These findings at least established that activation of mTORC1 was coincident with atrophy, and therefore was not sufficient to prevent muscle loss in sarcopenic conditions. We therefore asked whether counter-regulating this age-associated increase in mTORC1 signaling may perhaps be beneficial for skeletal muscle, and thus we treated aged rats for six weeks with a rapalog, RAD001, at a clinically relevant low dose (we reverse-translated the low dose that had been used in a human study) (16). Treatment with a similar low dose of the rapalog RAD001, although with a distinct dosing regimen (intermittent dosing) had recently been shown to delay age-related changes in the kidney (17).

We were surprised to see that skeletal muscle mass increased rather than decreased as a result of mTORC1 inhibition. This was not due to adverse events such as edema; muscles - particularly the TA, showed increased mass, and examination of individual myofibers showed a trend towards increased cross-sectional area; very small atrophic fibers that are found with age were in particular absent with rapalog treatment. With age, there is a dramatic increase in fibers with central nuclei - a sign of muscle undergoing degeneration followed by regeneration. Treatment with the rapalog for six weeks decreased the number of myofibers with central nuclei by almost half, which is a marker that there was less functional degeneration, requiring subsequent regeneration. In line with this, there were also signs that functional denervation occured with age; this impression was bolstered molecularly by the demonstration that gene markers associated with denervation, including MuSK and several of the acetyl choline receptor genes, were increased with age - consistent with a prior report (43). These denervation markers were each counter-regulated by the rapalog, indicating that rapalog treatment prevented functional denervation, providing an additional mechanism for preservation of muscle mass.

Rapalog treatment decreased mTORC1 signaling detected by de-phosphorylation of S6K1 and its downstream target rpS6. Coincident with this, mRNA levels of the putative atrophy marker MuRF1 were significantly reduced. In addition to MuRF1, the metallothionein MT1 was downregulated by mTORC1 inhibition. We had previously shown that this is a high-fidelity marker of atrophy, and knocking out the MT genes in mice causes muscle hypertrophy (45). This finding too is consistent with the increase in mass observed in the present study, and provides further mechanistic rationale. As for the senescence markers p16 and p21, they were elevated in aged muscle when compared to young muscle, and reversed towards the “younger level” following rapalog treatment, consistent with what had been shown previously in geriatric satellite cells (46, 53). This reversal of senescent markers suggest the possibility of improved satellite cell function necessary for muscle homeostasis, reflected by the positive morphological changes we observed in rapalog treated muscles.

In summary, mTORC1 signaling is hyper-activated in aged muscle, and this apparently is causal of sarcopenia, since inhibition of this signaling can increase muscle mass. The inhibition of denervation and senescence markers, and the subsequent decline in atrophy markers gives a therapeutic rationale for treating aged sarcopenic patients with an mTORC1 inhibitor.

## Materials and Methods

### Animal maintenance and RAD001 treatment

Male Sprague Dawley rats were obtained from Envigo (Indianapolis, IN), and housed at their facility under specific pathogen-free (SPF) conditions until the appropriate age. When transferred to our facility, rats continued to be maintained under SPF conditions, with regulated temperature and light cycles (22°C, 12-hour light/12-hour dark cycle: lights on at 0600hours/lights off at 1800 hours), and unrestricted access to food (2014 Teklad Global 14% Protein diet (Envigo)) and water. Animals were acclimated for a minimum of 4 weeks before being used for experiments. For age time course studies, rats ranging from 6 to 27-months (n = 6-8 animals/group) were fasted from 0600 hours to 1200 hours (during the light on cycle) before being anesthetized and euthanized for end of study analysis.. Gastrocnemius muscles were collected for molecular analysis. For other studies, RAD001 (Novartis) was prepared as a microemulsion pre-concentrate at 2% (w/w). Prior to dosing, it was diluted to a working concentration in water. Vehicle control consisted of microemulsion pre-concentrate (equivalent to a dose), diluted in water. At 22-months, animals were dosed daily for 6 weeks with either RAD001 or vehicle per os. In parallel, rats aged 7-months received vehicle as young adult controls. Four hours after the last dose of RAD001 or vehicle, rats were anesthetized with 3.5% isofluorane, and euthanized by exsanguination and thoracotomy. Gastrocnemius, soleus, plantaris and tibialis anterior (TA) muscles were collected and weighed; TA and plantaris muscles were processed as described below further assessment. All animal studies were done in accordance with institutional guidelines for the care and use of laboratory animals as approved by the Institutional Animal Care and Use Committee (IACUC) of the Novartis Institutes of Biomedical Research, Cambridge MA.

### Protein extraction and Western Blot analysis

Protein extracts were prepared from tibialis anterior muscles. In short, snap frozen tissue was pulverized in liquid nitrogen by mortar and pestle to a fine powder. Approximately 30mg of tissue powder was homogenized in MSD lysis buffer (#R60TX, Meso Scale Discovery,) supplemented with Protease and Phosphatase inhibitor cocktail (Thermo Fisher Scientific, MA). Following a 30 min incubation at 4°C with agitation, protein lysates containing the cytoplasmic fraction were collected via microcentrifugation. Protein concentration was determined by BCA protein assay (Thermo Fisher Scientific, MA), prior to Western blot analysis. Diluted proteins were separated by sodium dodecyl sulfate polyacrylamide gel electrophoresis (SDS-PAGE) on a 4-20% gradient Criterion TGX Precast Midi Protein gel (Bio-Rad, CA), and subsequently transferred onto nitrocellulose membranes (Bio-Rad, CA) with the Trans Turbo Blot system (Bio-Rad). Membranes were blocked in 5% milk in TBST for 1 h at room temperature, and incubated with primary antibodies overnight at 4°C. After three washes in TBST, membranes were incubated in the appropriate HRP conjugated secondary antibodies (Cell Signaling Technologies) for 1 h at room temperature. The following primary antibodies were used: anti-GAPDH (#5174, Cell Signaling Technologies, MA), anti-rpS6 (#2217), anti-p-rpS6(S240/244) (#2215), anti-S6K1 (#2708), anti-pS6K1(T389) (#9234), all from Cell Signaling Technologies. Anti-rabbit and anti-mouse IgG HRP-conjugated secondary antibodies were also from Cell Signaling Technologies. Densitometric analysis was performed using Fiji 1.51n software.

### Cryosectioning of frozen tissue

Plantaris muscles were removed, embedded in OCT (Tissue-Tek) and flash frozen in chilled 2-methylbutane (Fisher Scientific). Muscles were sectioned transversely with the Leica CM3050 S microtome and 10μm thick sections were collected for Hematoxylin and Eosin (H&E) staining and immunohistochemistry.

### Hematoxylin and Eosin staining

Muscle sections were fixed in 4% paraformaldehyde on ice for 10 mins and rinsed briefly with water five times. H&E staining was done using the Tissue-Tek Prisma automated slide stainer (Sakura Finetek). Images were captured using the Aperio ScanscopeAT (Leica Biosystems), and used to determine morphological changes, including the incidence of central nuclei.

### Immunohistochemistry

Myofiber cross-sectional area was measured on muscle cross-sections immuno-stained with the anti-Laminin antibody. Briefly, tissue sections were fixed in 4% paraformaldehyde on ice for 10 mins, and washed in 1X PBS prior to permeabilization in 0.3% Triton-X 100 in PBS. Non-specific sites were blocked in 16% goat serum diluted in 0.01% Triton-X 100 in PBS (blocking buffer) for 1 h at room temperature. Sections were incubated in anti-Laminin (Sigma Aldrich, L9393) antibody diluted at 1:1000 in blocking buffer overnight at 4°C. Primary antibody was detected by a 1h incubation with AlexaFluor-conjugated goat anti-rabbit secondary antibody (Life Technologies, A-11072) diluted in blocking buffer. Following a series of washes in 0.01% Triton-X in PBS, slides were mounted with Fluoromount-G (SouthernBiotech). Images were captured using the VS120 Virtual Slide Microscope (Olympus).

### RNA extraction, cDNA synthesis and Quantitative RT-PCR (RT-qPCR)

Tibialis anterior muscle was ground to powder as described above. Approximately 30mg of tissue powder was then processed using the miRNeasy Micro kit (Qiagen) according to the manufacturer’s protocol. RNA concentration was quantified by NanoDrop Spectrophotometer (NanoDrop Technologies), and quality was confirmed by the OD260/OD280 absorption ratio (>1.8). Following the manufacturer’s protocol, cDNA was synthesized from 1μg of RNA using the High Capacity cDNA Reverse Transcription kit (Applied Biosystems by Thermo Fisher Scientific). cDNA was diluted 1:10 in Ultra Pure Distilled RNase-free water (Invitrogen) prior to being used for further steps. The Standard TaqMan Gene Expression Master Mix (Applied Biosystems by Thermo Fisher Scientific) was used for all RT-qPCR reactions, and samples were run using a 384-well optical plate format. Reactions were performed using the ViiA 7 RT-qPCR System (Life Technologies), and data analyzed by the ΔΔcT method. TaqMan probes were optimized by Applied Biosystems: TATA-box binding protein (TBP) (Rn01455646_m1), Vps26a (Rn01433541_m1), MurF1 (Rn01639111_m1), MaFbx (Rn00591730_m1), Mt1 (Rn00821759_g1), Mt2A (Rn01536588_g1), Cdkn1a (Rn00589996_m1), Cdkn2a (Rn00580664_m1), Chrna1 (Rn01278033_m1), Chrne (Rn00567899_m1), MuSK (Rn00579211_m1), Myogenin (Rn00567418_m1) and Gadd45a (Rn01425130_g1).

### Statistical analysis

Statistical significance was determined by a one way ANOVA followed by Dunnett’s multiple comparison tests. Means from all groups were compared to the mean of the aged vehicle treated group, except where specified. All data are displayed as means with standard deviation. GraphPad Prism 7.04 software was used for calculations and graphing.

## Acknowledgements

The authors thank Danuta Lubicka for formulating RAD001, as well as study support associates and the veterinary team for maintaining aged rats, and assistance in animal experimentation. Thank you also to Bret Morin for collecting and processing tissues, and Paola Capodieci and Kristie Wetzel for assistance with histological imaging. We thank Samuel Cadena and the entire Age-Related Disorders group, and also the NIBR community for their enthusiastic support. All authors were employees of Novartis when this work was conducted, and some are current stockholders of Novartis.

---

**Figure S1.**
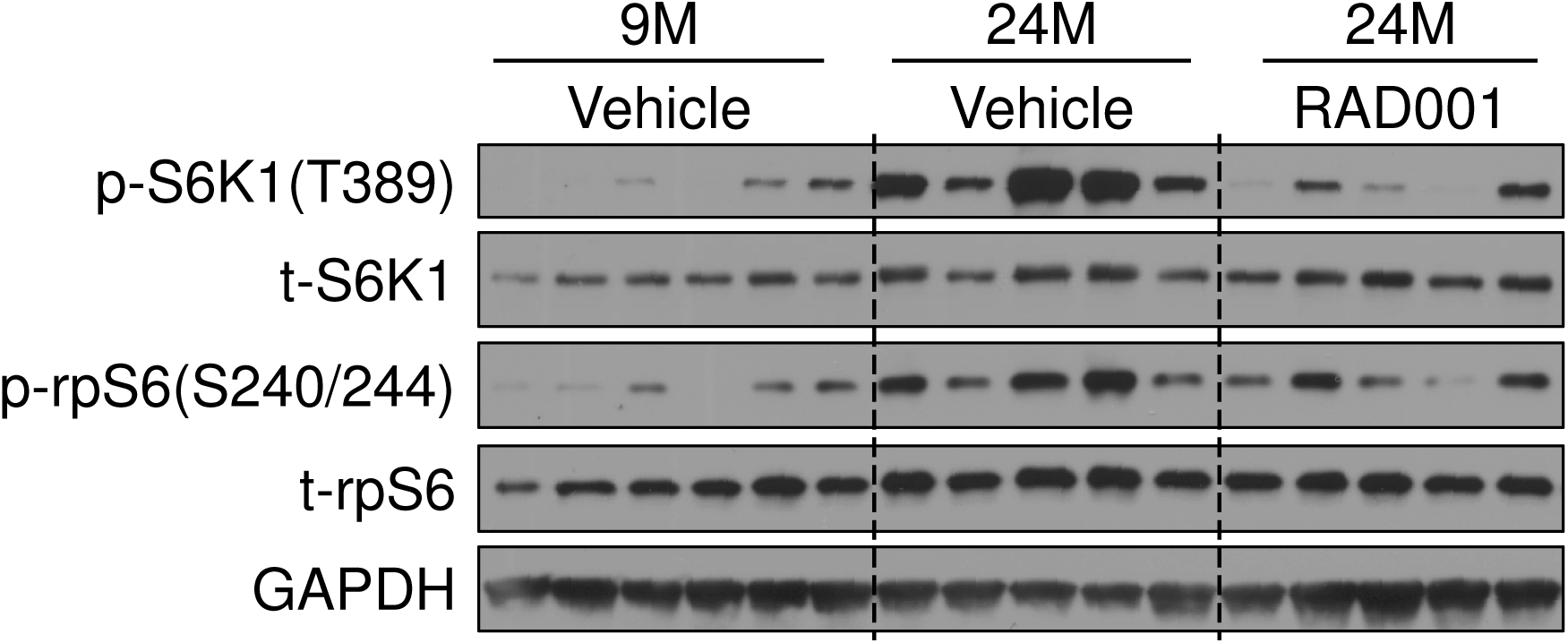
Rapalog treatment blocks mTORC1 pathway activity. Immunoblots of additional samples that were included in the analysis of phosphorylated (p) and total (t) protein for S6K1 and rpS6 in tibialis anterior muscles of 9- and 24-month old rats treated with vehicle and 24-month old rats treated with RAD001 in Fig. 3B, C. Glyceraldehyde-3-phosphate dehydrogenase (GAPDH) is shown as a loading control.

